# Clockor2: Inferring global and local strict molecular clocks using root-to-tip regression

**DOI:** 10.1101/2023.07.13.548947

**Authors:** Leo A. Featherstone, Andrew Rambaut, Sebastian Duchene, Wytamma Wirth

## Abstract

Molecular sequence data from rapidly evolving organisms are often sampled at different points in time. Sampling times can then be used for molecular clock calibration. The root-to-tip (RTT) regression is an essential tool to assess the degree to which the data behave in a clock-like fashion. Here, we introduce Clockor2, a client-side web application for conducting RTT regression. Clockor2 uniquely allows users to quickly fit local and global molecular clocks, thus handling the increasing complexity of genomic datasets that sample beyond the assumption homogeneous host populations. Clockor2 is efficient, handling trees of up to the order of 10^4^ tips, with significant speed increases compared to other RTT regression applications. Although clockor2 is written as a web application, all data processing happens on the client-side, meaning that data never leaves the user’s computer. Clockor2 is freely available at https://clockor2.github.io/.

## Introduction

Phylodynamic analyses make use of genetic sequence data to understand the evolution, epidemiological, and ecological dynamics of a pathogen. Importantly, phyodynamics achieves its greatest value when generating insight about infectious disease dynamics outside the purview of epidemiology. This frequently occurs at population interfaces, such as during transmission across host sub-populations, geographical boundaries or host species. Despite the increasing complexity of such datasets, the essential component to all phylodynamic modelling is the assumption of a molecular clock relating epidemiological and evolutionary timescales (Biek et al., 2015).

The simplest molecular clock model is the strict clock, which assumes a constant rate of substitution per unit time, known as the ‘evolutionary rate’. When the evolutionary rate is constant throughout a phylogenetic tree, the term global molecular clock is used. In contrast, a strict local clock refers to the situation where different substitution rates apply to different monophyletic groups within a tree (Ho and Duchêne, 2014). The branches of local clocks are sometimes referred to as ‘foreground’ while the remaining branches are known as the ‘background’ (Yoder and Yang, 2000). The assumption of a local clock may, for example, correspond to sampling from different host populations, host species, or pathogen lineages (Worobey et al., 2014). Clockor2 includes examples from Dudas et al. (2018) where local clocks are fit to MERS-CoV samples from human and camel hosts, and another from Porter et al. (2023) with SARS-CoV-2 samples from human and mink hosts. Clockor2 enables rapid inference of global and local strict molecular clocks from phylogenetic trees where the tips are annotated with sampling times and other relevant data, using root-to-tip regression (RTT). Several other tools allow for the inference of strict molecular clocks via RTT, but none readily offer the ability to fit local clocks models (Rambaut et al., 2016; Hadfield et al., 2018; Sagulenko et al., 2018; Volz and Frost, 2017).

Phylodynamic datasets are and will continue to grow in size and scope Featherstone et al. (2022). For example, datasets of thousands to tens-of-thousands of samples have been used to understand the spread of SARS-CoV-2 at international scales, the emergence of variants of concern (VOC), and transmission between species (du Plessis et al., 2021; Hill et al., 2022; Nadeau et al., 2023; Porter et al., 2023). However larger datasets, as a function of their size, are more likely to sample from distinct populations, making local clocks increasingly important. Currently, testing the fit of a local clock over alternative models, such as global or relaxed clocks, frequently requires intensive computational efforts using common Bayesian phylodynamic applications such as BEAST or RevBayes (Bouckaert et al., 2019; Suchard et al., 2018; Höhna et al., 2016; Drummond and Rambaut, 2007; Drummond et al., 2012), each requiring hours to days of computational time. Clockor2 uniquely offers a scalable and accessible client-side web application for exploring the fit of local clocks, with results available in seconds to minutes to direct subsequent phylodynamic analysis.

Specifically, Clockor2 allows users to perform RTT regression for fitting global and local clocks (Fig 1). The user begins by dropping or importing a newick tree. Sampling dates and group identifiers can then be parsed from tip labels or separate files on input. Like other RTT regression applications, Clockor2 also allows users to infer the best fitting root based on the *R*^2^ value or residual mean square (RMS) of the RTT regression. Both are key indicators of clock-like evolution. It also offers users a local clock-search function to test assumptions about the number of local clocks in a dataset as well as the ability to add a local clock interactively.

**Figure 1:**
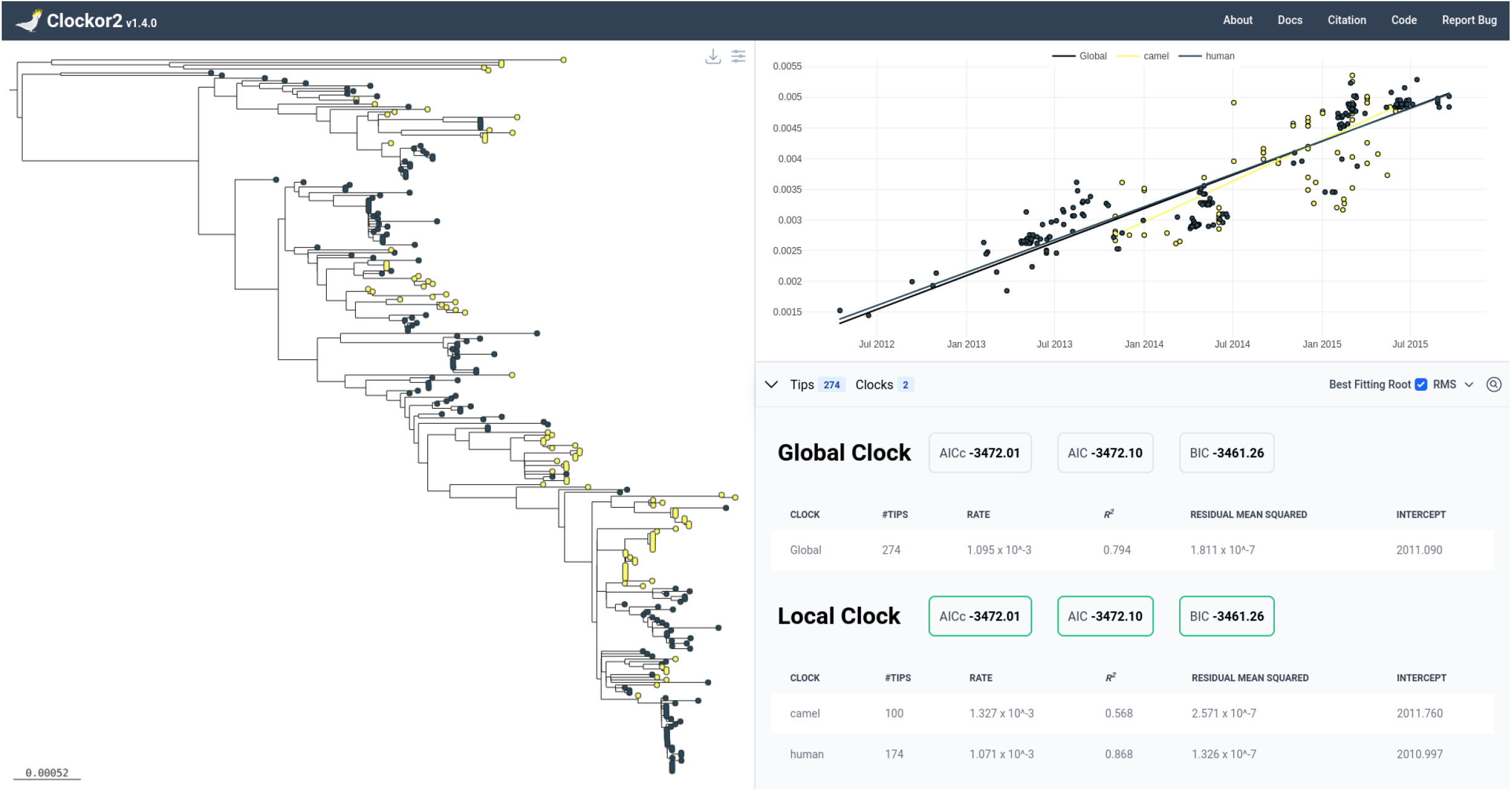
Clockor2 presents the tree alongside RTT regression data. Users can toggle between local and global clocks and alter the appearance of the tree.

## Methods

### General model for global and local strict clocks

RTT regression consists in modelling the evolutionary rate as the slope of a linear regression of the distance from the root to each tip (RTT distance), typically in units of substitutions per site (subs/site), against the sampling date of each tip (Drummond et al., 2003). If we denote the evolutionary rate as *r* (usually in units of *subs/site/time*), RTT distance as *d* (usually in units of *subs/site*), *o* as the intercept (interpreted as origin), and sampling times as *t*, then the model for a global strict clock takes the form:

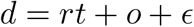

where *ϵ* is an error term.

Clockor2 uses a generalisation of this model to accommodate local clocks. For a given tree with a set of tips *T*, we define local clocks as pertaining to *groups* of tips *g*_*i*_ and a rate parameter for each (*r*_*i*_). For a strict clock model with two local clocks, we then write:

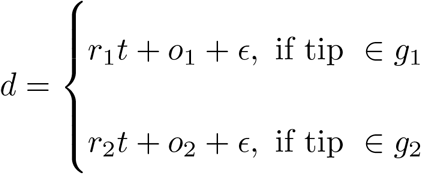

We refer to *groups* instead of *clades* because while collections of tips belonging to one local clock necessarily share a common ancestor, they do not necessarily comprise a whole clade. This occurs when two or more local clocks are nested. The tips comprising the outer clock(s) cannot comprise a whole clade if another local clock is nested within it. For example, local clock 1 in Fig. 2 B,D does not comprise a clade (i.e. is not monophyletic) because local clock 2 is nested within it.

**Figure 2:**
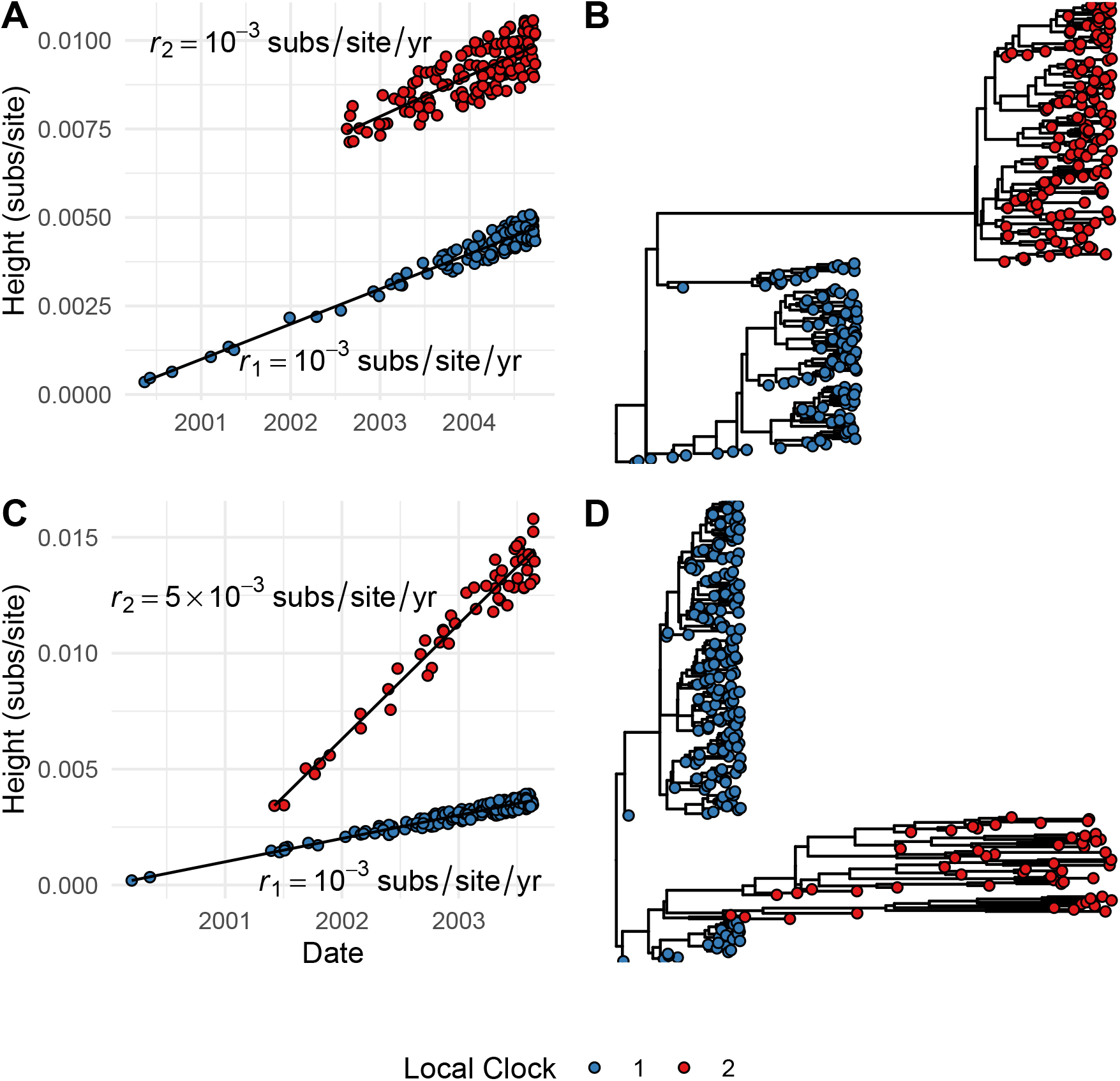
Simulated examples of how local clocks may manifest in trees and RTT regression data. (**A**) RTT regression data for two local clocks with similar rates separated by a long branch. (**B**) A tree characteristic of two similar local clock rates separated by a long branch. (**C**) RTT regression data where two local clocks have different evolutionary rates. (**D**) A tree characteristic of two local clocks with different rates.

This general model then captures the two key scenarios where local clocks may be appropriate. The first is where rates are similar between local clocks, but separated by a long branch (Fig. 2A-B). For example, in the evolution of VOCs in SARS-CoV-2 or due to temporally-sparse sampling in the case of ancient *Yersisnia pestis* samples (Tay et al., 2022; Eaton et al., 2023; Hill et al., 2022).The second scenario is where rates differ between local clocks (Fig 2 C-D). For example, this can occur when a pathogen spreads in different host species, such as has been observed for SARS-CoV-2 in mink and human hosts (Porter et al., 2023).

For each group of tips defining a local clock, we independently conduct a RTT regression to estimate the evolutionary rate (slope). *R*^2^ or RMS values are then an indication of clock-like behaviour for each local clock. Clockor2 focuses on *R*^2^ and RMS as indicators of clocklike evolution the because they offer offer the most straightforward interpretation of clock-like evolution. *R*^2^ values of one indicate perfect clock like evolution, while values of zero indicate a lack of a molecular clock. Likewise, lower RMS values indicate better fit of a strict clock.

Local clock and or global clock configurations can also be compared using an information criterion that combines the likelihood of each local clock’s RTT regression while penalising the number of inferred parameters (three for each clock slope, intercept, and variance). Clockor2 allows users to use either the Bayesian Information Cirterion (BIC), Aikake Information Criterion (AIC), or corrected Aikake Information Cirterion (AICc). We recommend using the BIC because it most heavily penalises the addition of extra parameters, and local clocks in turn.

Derivations of the above information criteria for the local clock model are given in the supplementary methods. Briefly, these exploit the assumption of independent sampling to factor the likelihood across local clocks. Note however that this assumption is always flawed because samples necessarily share some ancestry by the assumption of a phylogenetic tree. In other words, ancestral branches are counted over many times in the calculating the distance from root to tip for each sample. However, this is a limitation of the RTT regression approach generally, rather than of Clockor2 itself.

### Algorithm for local clock search

Where it is hypothesised that a datasset contains local clocks, Clockor2 provides functionality to corroborate the this hypothesis by performing a search for local clocks in the tree. Briefly, the algorithm takes a maximum number of clocks and a minimum number of tips (group size) for each local clock as input parameters. It then iterates through all combinations of internal nodes from which local clocks can descend to induce corresponding local clock configurations. Importantly, the clock search algorithm tests for a number of clocks up to and including the maximum number so that more parsimonious configurations with fewer clocks may be found. Configurations are compared using the information criteria outlined above. Again, we recommend the BIC as it penalises additional parameters (ie. additional local clocks) most heavily. See here for an animation of the clock search algorithm.

We stress that this algorithm is intended to corroborate hypotheses about a particular local clock configuration, but is not intended to be performed as a blind search for local clocks. This is because it is highly prone to over-fitting where the maximum number of clocks is inflated, as outlined later in the results section.

The clock-search algorithm operates in polynomial time (see supplementary material). Efficiency is improved by reducing the maximum number of local clocks in the search, increasing the minimum group size, and contingent on the topology of the underlying tree. However, the former two parameters exert a far greater effect on efficiency than topology.

### Clock-search algorithm simulation study

We conducted a simulation study to test the accuracy of the clock-search algorithm. We started with a core set of 100 simulated trees of 250 tips and added a local clock descending from a randomly selected node such that it would contain between 50 and 150 tips. For each, we simulated a 5-fold rate increase occurring in either the stem branch leading to the group/clade, or throughout the group. These scenarios are characterised in Fig 2 C,D respectively. For each of the resulting 200 trees, we applied the clock search algorithm with a minimum group size of 50 tips and a maximum number of of 2-5 clocks. The case of a maximum of 2 clocks tests for baseline accuracy with the search parameters match reality. Searches with a maximum of 3-5 clocks test for over-sensitivity in the algorithm where the maximum number of clocks is inflated. All clock-search tests use the BIC.

### Finding the Best Fitting Root

Clockor2 selects the best fitting root based on the *R*^2^ of RMS of a global clock model for the input tree. It follows the same algorithm as implemented in Tempest (Rambaut et al., 2016), but makes use of parallelisation to improve speed for larger trees. Briefly, the tree is rooted along the branch leading to each internal node or tip, an RTT regression is performed, and the root position along the branch leading to the highest *R*^2^ or RMS is selected. When targeting *R*^2^, Clockor2 optimises the root position using the golden-section search algorithm (Kiefer, 1953). There is an analytical solution for the RMS (See supplementary section 3).

The best fitting root is inferred using a single, global clock because it presents the most parsimonious model of the evolutionary rate for a given tree. The fit of more elaborate local clock models can then be compared to this using information criteria and/or comparing the *R*^2^ or RMS values of each model. Clockor2 does not find the best fitting root for local clock models because the search space of best fitting roots and local clock configurations quickly becomes prohibitive and is possibly unidentifiable.

### Dependencies

Clockor2 has three key dependencies for handling, and plotting trees and RTT data. Trees are handled and manipulated using the phylojs library. Phylocanvas is used to viaualise trees and plotly.js is used to plot RTT data (Abudahab et al., 2021; Plotly-Technologies-Inc., 2015).

## Results

### Efficiency

Clockor2 can process trees of up to the order of 10^4^ tips, thus fit for the expanding size and diversity of phylodynamic datasets. Finding the best fitting root makes use of parallelisation to increase speed. Speedup is therefore proportional to the number of threads or cores available, in addition to the choice of browser and computer. For example, Clockor2 is faster than TempEst v 1.5.3 a 2021 Macbook pro with 16Gb of memory and 8 cores running chrome v113.0.5627.126, (Table 1). However, we found Clockor2 to be comparable or slower on other combinations of processor and browser.

**Table 1:** Time in seconds taken to find the best fitting root for test trees of 100, 500, 1000, 5000, and 10000 tips using Clockor2 and TempEst v1.5.3 on a 2021 Macbook with 16Gb of memory and 8 cores running chrome v113.0.5627.126. Times vary with computer and browser. In general the relative efficiency of Clockor2 will increase with the number of cores. Using the residual mean squared as an optimisation target is faster because there is an analytical solution.

| Tips | $R^2$ | | Residual Mean Squared | |
| --- | --- | --- | --- | --- |
|  | Clockor2 | TempEst | Clockor2 | TempEst |
| 100 | 0.313 | 0.760 | 0.129 | 0.050 |
| 500 | 1.370 | 2.400 | 0.502 | 1.500 |
| 1000 | 3.476 | 10.280 | 1.514 | 5.430 |
| 5000 | 78.013 | 272.290 | 24.370 | 122.000 |
| 10000 | 306.821 | 1310.340 | 94.992 | 951.000 |

The user interface also remains responsive when working with large trees. This is in large part due to the use of WebGL in the tree and plotting components, which exploit GPU acceleration to render large and interactive trees and datasets through Phylocanvas and Plotly.js respectively (Abudahab et al., 2021; Plotly-Technologies-Inc., 2015).

### clock-search accuracy

For clock searches with a maximum number of 2 clocks, the clock-search algorithm correctly identified 2 local clocks in the simulated data with 100 percent accuracy (Fig. 3). However, when the algorithm was allowed to search for configurations with 3-5 clocks, only 13 and 17 analyses correctly recovered 2 clocks for the stem only and stem+clade rate increases respectively. Although we used the BIC, the most conservative information metric used in Clockor2, the clock-search algorithm is clearly still highly prone to over-fitting. This is because a higher number of clocks allows for tighter clusters of points in the RTT regression to be found, which are favoured over a lower number of clocks.

**Figure 3:**
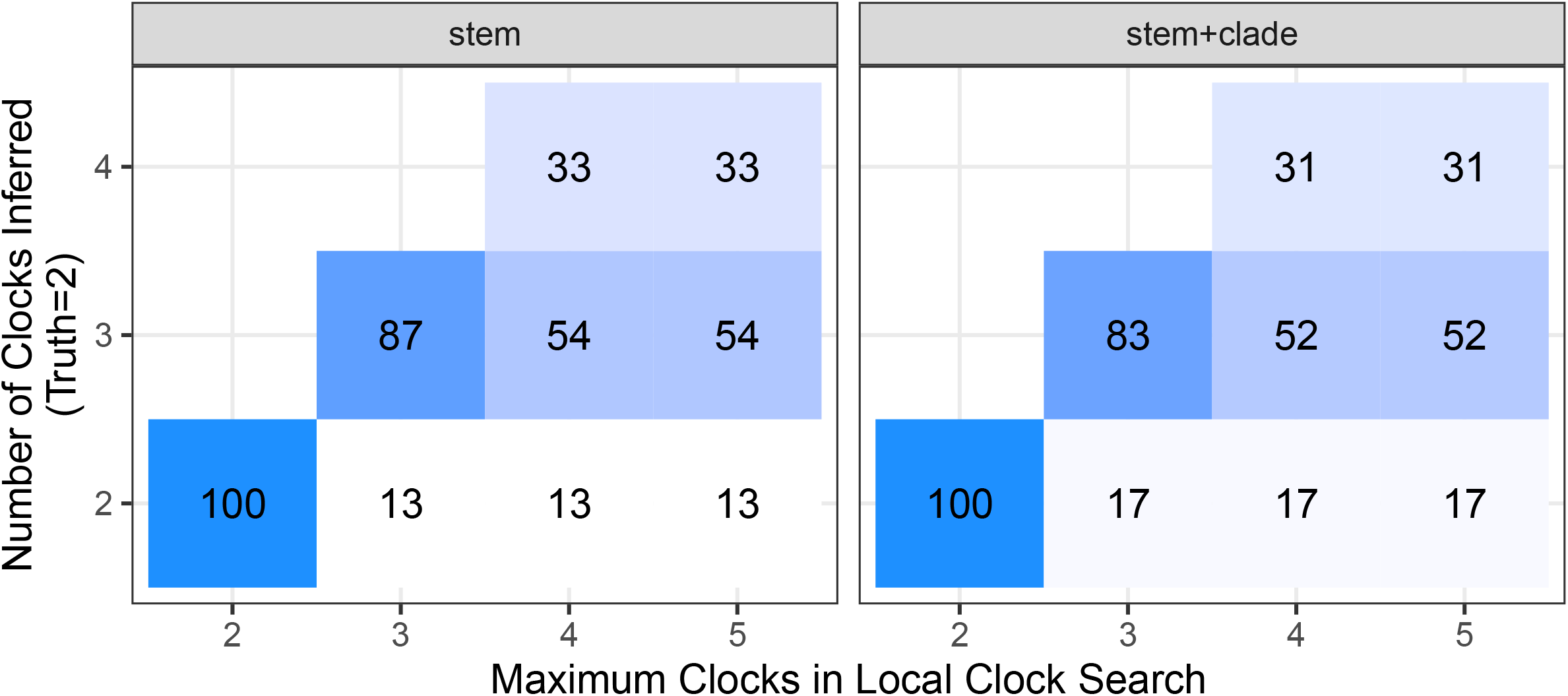
Confusion matrices comparing the number of inferred clocks against the maximum number of clocks allowed by each search, with a true number of 2 clocks in each search. “stem” and “stem+clade” refer to either a rate increase along only the stem of a clade, or with the rate increase continuing in the clade. The true value is 2 and accuracy decreases as the maximum number of clocks allowed by the clock-search algorithm increases. The colour of and number in each tile reflect the number of analyess out of 100 selecting the corresponding number of clocks.

To this end, we again emphasise that the clock-search algorithm is only intended to be used as a tool testing a number of clocks up to and including the hypothesised number, but never more. It should never be used to blindly search for local clocks in the absence of a biological hypothesis. The Clockor2 how-to documents provide example of over-fitting using the clock search function when applied to various empirical datasets (Porter et al., 2023; Dudas et al., 2018).

## Discussion

Clockor2 provides a flexible and scalable front-end web based RTT regression platform. Its extension to fitting local clocks allows it to accommodate the growing complexity of phylodyanmic datasets as genomic epidemiology plays a growing role in infectious disease surveillance.

As a front end application, Clockor2 is also highly accessible with no installation steps required, although users have option of saving the site to run locally. Wherever there is a browser, it is possible to conduct an RTT regression using Clockor2 with the data never leaving the user’s computer.

## Supporting information

Derivations for Information criteria, best fitting root analytical solution, and time complexity for clock search.

## Future Directions

In the future it will be possible to re-implement core functionality in increasingly popular and highly efficient programming languages, that can compile to Web-Assembly format. For example, as the bioinformatics ecosystem in Rust continues to develop, it will be possible to further improve the efficiency of Clockor2 using packages such as Bio-Rust (Köster, 2015).

## Data availability

All code required to replicate the simulation study and figures in the paper is available at https://github.com/LeoFeatherstone/clockor2Paper. The code for Clockor2 is open source at https://github.com/clockor2/clockor2.

## Acknowledgements

We are grateful to Luiz Max Carvalho for helpful correspondence in ascertaining the analytical solution for the position of the best fitting root using the root mean squared.

