## Supplementary material for "Clockor2: Inferring global and local strict molecular clocks using root-to-tip regression": Derivations for Information criteria, best fitting root analytical solution, and time complexity for clock search.

### 1 Information Criteria for Local Clocks

Clockor2 uses the AIC, AICc, and BIC to compare local clock models. Here we begin by giving the formula for each. Each criterion depends on the number of samples ($n$), number of inferred parameters ($k = 3$ per clock), and the likelihood of the model fit ($\hat{L}$). We capitalise upon the assumption of independant sampling in root-to-tip regression to factorise the likelihood and calculate information criteria for local clock models.

$$\text{BIC} = k \ln n - 2 \ln \hat{L} \tag{1}$$

$$\text{AIC} = 2k - 2 \ln \hat{L} \tag{2}$$

$$\text{AICc} = \frac{2k^2 + 2k}{n - k - 1} + k \ln n - 2 \ln \hat{L} \tag{3}$$

We begin with the example of two local clocks before generalising to an arbitrary number of clocks. Consider a tree with $n$ tips and 2 local clocks. Each local clock is comprised of a group of tips $g_1$ and $g_2$, such that $\|g_1\| + \|g_2\| = n$

The log-likelihood, containing the product of probability densities for each data point, can then be factored into the sum of log likelihoods for each local clock:

$$\ln \hat{L} = \sum_{i=1}^{2} \ln \hat{L_{g_i}} \tag{4}$$

The number of inferred parameters is also $k = 6 = 2 \cdot 3$ (intercept, variance, slope).

For example, we can then write the BIC as:

$$\text{BIC} = 6 \ln n - 2 \sum_{i=1}^{2} \ln \hat{L_{g_i}} \tag{5}$$

Generalising to the case where we have $c$ local clocks, the BIC then becomes:

$$\text{BIC} = (3c) \ln n - 2 \sum_{i=1}^{c} \ln \hat{L_{g_i}} \tag{6}$$

#### 2 Efficiency of clock search algorithm

We briefly provide reasoning for why the clock search algorithm operates in polynomial time. Consider a tree with $n$ tips, and a clock search that searches for a maximum of $c$ clocks each of a max size $s$, ensuring that $cs \leq n$.

Given there are $n$ tips, there are then $2(n-1)$ internal nodes from which a clock can descent, and for a given number of clocks $1, ..., c$, there are $p$ internal nodes such that a local clock can descend from each and induce a group with more than $s$ tips ($\|g\| \geq s$). $p$ depends on the topology of the tree (denoted as $t$) as well as the number of clocks ($c$) and minimum group size ($s$), so we now write $p(t, c, s)$ with $1 \leq p(t, c, s) \leq n$.

The number of possible combinations for of nodes that can be chosen to fit $c$ local clocks, each of at least size $s$ is then:

$$\binom{p(t,c,s)}{c} = \frac{p(t,c,s)!}{c!(p(t,c,s)-c)!} \tag{7}$$

That is, the clock search is polynomial in $p(t, c, s)$. $p(t, c, s)$ is related to $n$ by the topology of the particular tree being searched, so we assert for generality that the clock search is polynomial in $n$. $p(t, c, s) = n$ when $c = s = n$.

Finally, the clock search searches for a $1, ..., c$ clocks, completing in a number of steps equal to:

$$\sum_{1}^{c} \binom{p(t,c,s)}{i} \tag{8}$$

The number of steps can be reduced by reducing the number of clocks searched for and increasing the minimum group size.

As noted above, tree topology also affects $p(t, c, s)$ also affects the number of steps taken to complete the clock search, but we expect this to affect efficiency the least out of the tree parameters, and is usually expected to be beyond the control of the researcher anyway. For completeness, we note that a perfectly balanced tree requires the most steps for the algorithm to complete while a completely imbalanced, ladder tree would require the least.

#### 3 Best Fitting root Targeting Residual Mean Squared

When searching for the best fitting root, for each proposed root there is the question of where to place the root between the two descending branches to maximise temporal signal. In Clockor2, this point is selected by maximising $R^2$, or minimising the residual mean squared. For the residual mean square, there is an analytical solution for this point that we derive here.

The following is an independent derivation by LAF of the algorithm implemented in Tempest and derived by Luiz Max Carvalho.

First, define the following:

1. $L$ is the sum of basal branch lengths leading to the left and right root-child.
2. $d_i$, The set of root-to-tip distances where all basal branch length ($L$) is shifted to the branch leading to the right root-child.
3. $t_i$, The sampling times associated with each $d_i$
4. $c_i$, An indicator variable indicating whether a tip descending from the left or right root-child. $c_i = -1$ if descending from the left child, and $c_i = 1$ if descending from the right.
5. $x$ is the portion of total branch length $L$ that we add to $d_i$ descending from the left root-child and subtract from $d_i$ descending from the right root-child

Now, to capture the different to root-to-tip distances that come with shifting the root position, we define:

$$d_i' = d_i + c_i x \tag{9}$$

And we seek to minimise with respect to $x$ the Residual Mean Square:

$$\mathrm{RSM}(x) = \frac{1}{n-2} \sum_i (\hat{d}_i' - d_i')^2 \tag{10}$$

Which is equivalent to minimising the Residual Sum of Squares(RSS)

$$\begin{aligned}\text{RSS}(x) &= \sum_i (\hat{d}'_i - d'_i)^2 \\ &= S_{d'd'} - S_{td'}^2 S_{tt}^{-1}\end{aligned} \tag{11}$$

$$\text{Where } S_{d'd'} = \sum_i (\bar{d}' - d'_i)^2, S_{tt} = \sum_i (\bar{t} - t_i)^2, \text{ and } S_{d't} = \sum_i (\bar{d}' - d_i)(\bar{t} - t_i) \tag{12}$$

Subbing $d'_i = d_i + c_i x$ into each of $S_{d'd'}$, $S_{tt}$, and $S_{d't}$, we come to:

$$\text{RSS}(x) = S_{dd} - 2xS_{dc} + x^2 S_{cc} - S_{tt}^{-1}(S_{dt} - 2xS_{dt}S_{tc} + x^2 S_{tc}^2) \tag{13}$$

Observe that RSS$(x)$ is quadratic in $x$, and that RSS $\geq 0$, so there must be a global minimum we can solve for.

We therefore solve:

$$\begin{aligned}&\frac{d}{dx}\text{RSS}(x) = 0 \\ &\implies -2S_{dc} + 2xS_{cc} + 2S_{td}S_{tc}S_{tt}^{-1} - 2xS_{tc}^2 S_{tt}^{-1} = 0 \\ &\implies x = \frac{S_{dc} - S_{ty}S_{tc}S_{tt}^{-1}}{S_{cc} - S_{tc}^2 S_{tt}^{-1}}\end{aligned} \tag{14}$$

Finally, it may be that $x > L$. For this reason, Clockor2 internally relies on $\alpha = \frac{x}{L}$, the proportion of $L$ assigned to the left branch under the best fitting root. Then, we take the following as the solution:

$$\alpha = \max\left\{\min\left\{\frac{S_{dc} - S_{td}S_{tc}S_{tt}^{-1}}{L(S_{cc} - S_{tc}^2 S_{tt}^{-1})}\right\}, 1), 0\right\} \tag{15}$$
